# Repeated Subconcussive Head Impacts Compromise White Matter Integrity and Bimanual Coordination in Collegiate Football Players

**DOI:** 10.1101/2025.05.14.654080

**Authors:** Sebastian D’Amario, Blaire Magee, Allen A. Champagne, Madison E. Wilson, Gabriel Ramírez-García, Nicole S. Coverdale, Douglas J Cook

## Abstract

**Objectives:** To determine whether repetitive subconcussive impacts in collegiate football are associated with altered white matter microstructure, motor control deficits, and changes in concussion symptoms across a single season.

**Design:** Cohort study with non-contact controls and stratification of contact athletes by head-impact exposure.

**Method:** Twenty-two male varsity football players and 27 non-contact controls underwent pre-season diffusion tensor imaging. Tract-specific fractional anisotropy (FA) and mean diffusivity (MD) were derived from commissural and association tracts using probabilistic tractography. Helmet-mounted accelerometers quantified head-impact frequency and gForce over the season, classifying athletes into high (HE) and low-exposure (LE) groups. Football players completed the Kinarm Ball-on-Bar bimanual coordination task and SCAT3 symptom checklist pre- and post-season.

**Results:** At pre-season, contact athletes showed altered white matter microstructure versus controls, with higher FA and predominantly lower MD across most tracts. Over the season, HE athletes sustained more total impacts and gForce than LE athletes and showed declining bimanual coordination on the Ball-on-Bar motor task. SCAT3 symptom and severity scores were low and showed no differences in change.

**Conclusions:** Repetitive subconcussive exposure in collegiate football is associated with persisting white matter differences and subtle bimanual motor coordination deficits that are not detected by routine symptom-based tools, supporting the use of advanced neuroimaging and robotic motor assessment to monitor athlete brain health.

## Introduction

Head injuries sustained during contact sport participation are increasingly recognized for their potential long-term effects on brain health.^1–3^ While concussions have been extensively studied, increasing attention has focused on the effect of repetitive subconcussive impact (RSCI) exposure, as they occur much more frequently than concussion. In particular, high school and collegiate football players experience an average of 600 to 1400 impacts per season, depending on the level of play and position.^1,4^ A subconcussive impact is defined as a head impact that does not produce observable clinical symptoms yet may still result in neurophysiological changes,^1,2^ making them challenging to detect and address in real time. Nonetheless, neuroimaging studies have highlighted measurable changes in brain structure, particularly in WM microstructure. These changes may occur because the biomechanical forces underlying RSCIs (acceleration-deceleration impacts) may result in axonal shearing and neuronal disruption, initiating a cascade of neurochemical events that compromise WM integrity.^3,5^

Diffusion tensor imaging (DTI) has been instrumental in quantifying changes in WM integrity with metrics such as fractional anisotropy (FA) and mean diffusivity (MD).^6,7^ FA measures how directionally restricted water movement is, while MD captures the overall amount of diffusion. Alterations in these metrics are associated with microstructural damage processes, including axonal injury and demyelination.^8^ However, the findings in RSCI-related DTI studies have often been inconsistent, highlighting heterogeneity in WM changes.^9,10^ For instance, FA and MD have both been shown to decrease^11–14^ and increase^14,15^ after RSCIs. Given these inconsistencies in DTI findings, a multimodal approach to capture holistic changes from RSCI is very valuable.

Given that RSCIs produce measurable biomechanical strain,^4,16^ it is reasonable to expect that these effects extend to players’ motor behaviour. The Kinarm robotic assessment system provides standardized tasks, supported by large normative datasets, that can detect subtle changes in motor performance. In particular, the Ball-on-Bar (BOB) task is a validated measure of bimanual coordination^17^ and has been used to identify clinical consequences in stroke^17^ and amyotrophic lateral sclerosis (ALS).^18^ As individuals with long-term exposure to RSCI frequently report headaches, cognitive slowing, noise sensitivity, short-term memory problems, and disrupted sleep,^19–21^ we also included the Sport Concussion Assessment Tool, third edition (SCAT3), to capture subjective symptom burden and severity.^22^ Moreover, combining these behavioural and clinical measures with neuroimaging provides a more comprehensive view of the subtle and cumulative effects of RSCI on white matter integrity.^11,19,23^

In this study, our aim was to examine white matter microstructure in Canadian Varsity collegiate football players over the course of a football season. By stratifying players into high- and low-exposure groups using helmet accelerometer data, we examined how cumulative RSCIs affect motor performance and clinical scoring, and how these effects change over the course of a football season. We hypothesized that higher RSCI exposure would be associated with measurable FA and MD changes, reflecting white matter microstructural compromise. These changes are expected to be found in tandem with speed and accuracy deficits in players with greater RSCI exposure compared to those with low exposure. These findings may contribute to a more comprehensive understanding of the neuropathophysiology of RSCIs to formulate future strategies for mitigating risks in contact sports.

## Methods

### Participants

Twenty-two male athletes from the Queen’s University Varsity Football Team (20 ± 1 years old) in Canada, along with 27 controls (21 ± 2.5 years) participated in this study. Neuroimaging, behavioural and neuropsychological assessments (SCAT3) were conducted at two time points for the subconcussed group: PRE and POST (30 days following season’s end). All athletes provided written informed consent, and the study was approved by the Queen’s University Health Sciences Research Ethics Board. Participants all played in at least one previous season, and were excluded if they had pre-existing injuries, sustained injuries during the season that affected their ability to participate or had been diagnosed with a concussion within the past year or during the season.

### Impact Monitoring

Subconcussive impact exposure was quantified using gForce Tracker (GFT) accelerometers (Artaflex Inc., Markham, Ontario, Canada). The devices were mounted inside each athlete’s helmet using 3M Dual Lock™ Velcro, positioned at the left crown. Each accelerometer, capable of measuring linear acceleration and rotational velocity, was configured with a 15g threshold to exclude non-relevant movements (e.g. running, setting helmet down). During games and practices, on-site spotters flagged non-head impact events (e.g., dropped helmets) for subsequent removal. After a manual review of these instances, we calculated the session average and cumulative impacts. Athletes were then stratified into high (HE; n = 11) and low (LE; = 11) exposure groups using a median average frequency of impacts per session^24^ of 10 as the threshold.

### MRI Acquisition

All MRI scans were conducted using a Siemens 3.0 T Magnetom Tim Trio system. T1-weighted structural images were acquired using an MP-RAGE sequence (TR = 1760 ms; TE = 2.2 ms; TI = 900 ms; matrix = 256 x 256; FOV = 256 mm^2^; voxel size = 1 x 1 x 1 mm). We performed a diffusion-weighted fast spin-echo (TSE-DWI) sequence with the following parameters: TR = 7800 ms; TE = 95 ms; FOV = 256 mm^2^; matrix = 128 x 128; 60 axial slices; voxel size = 1 x 1 x 1 mm, b-value = 1000 s/mm^2^; and 30 gradient directions. Additionally, three non-diffusion-weighted (b = 0) images were acquired for normalization. Images were collected at the three time points under consistent conditions. Additionally, the SCAT3 symptom checklist was self-reported prior to each MRI. Higher scores on the SCAT3 indicate that the presence and severity of symptoms are more severe.

### Neuroimaging Processing

Diffusion data were preprocessed using tools from FMRIB’s Software Library (FSL).^25^ Preprocessing steps included correction for susceptibility-induced distortions using the *topup* tool, correction for eddy currents and motion artifacts using *eddy*, and brain extraction with the *bet* tool. Diffusion tensors were calculated for each voxel using *dtifit* to produce maps of fractional FA and MD metrics for each subject. These maps were aligned to a standardized Montreal Neurological Institute (MNI152 2 mm) template using *FLIRT* and *FNIRT* for linear and non-linear registration, respectively.

### Tractography

Following preprocessing, probabilistic tractography was conducted to analyze WM tracts using BEDPOSTX, which models crossing fibres within each brain voxel. ^26^ Streamline density maps, generated using FSL’s PROBTRACKX tool, quantified connectivity from the seed region to the target region for each tract. To ensure comparability across subjects, density maps were normalized by dividing each map by its waytotal using FSL’s *fslmaths*. Waytotals represent the total number of streamlines reaching the target mask from the seed mask, accounting for inter-subject differences in scan quality or motion. In total, this was done for the following tracts: Anterior Thalamic Radiation, Corticospinal Tract, Forceps Major, Forceps Minor, Inferior Fronto-occipital Fasciculus, Inferior Longitudinal Fasciculus, Superior Longitudinal Fasciculus, Superior Thalamic Radiation, Uncinate Fasciculus. Normalized tracts were registered to the MNI152 2mm standard space using *FLIRT* and *FNIRT* tools. Each tract was thresholded to eliminate low-probability streamlines, applying a consistent threshold of < 1% across subjects to ensure uniformity. Thresholded maps were then binarized to create masks for individual WM tracts. Tract-specific metric values were calculated by overlaying the binary masks onto standardized maps for each metric. The mean metric values for each subject and tract were extracted using FSL’s *fslstats* tool, enabling quantitative analysis of WM microstructure. Anatomical identification was done using the Human Xtract atlas.^27^

### Kinarm Task

Our assessment of motor ability had participants perform a bimanual BOB coordination task using the KINARM robotic interface that displayed a 30 cm virtual bar (1 cm thickness) connecting their index fingers and supported a 1 cm-radius virtual ball at its centre (Figure 1B). A gravitational constant (bar mass: 0.166 kg; ball mass: 0.4 kg) created realistic weight cues, and vision of the arms and ball was occluded.^17^ Participants were instructed to move the ball “quickly and accurately” to a sequence of four circular targets (1 cm radius) arranged 10 cm from the workspace origin, with each target turning from red to yellow once the ball entered and was held for 1 s. The task comprised three one-minute levels of increasing difficulty: Level 1 fixed the ball to the bar; Level 2 allowed the ball to translate along the bar and fall off at ≥ 20° tilt; and Level 3 increased the ball’s frictionless rolling speed.

**Figure 1.**
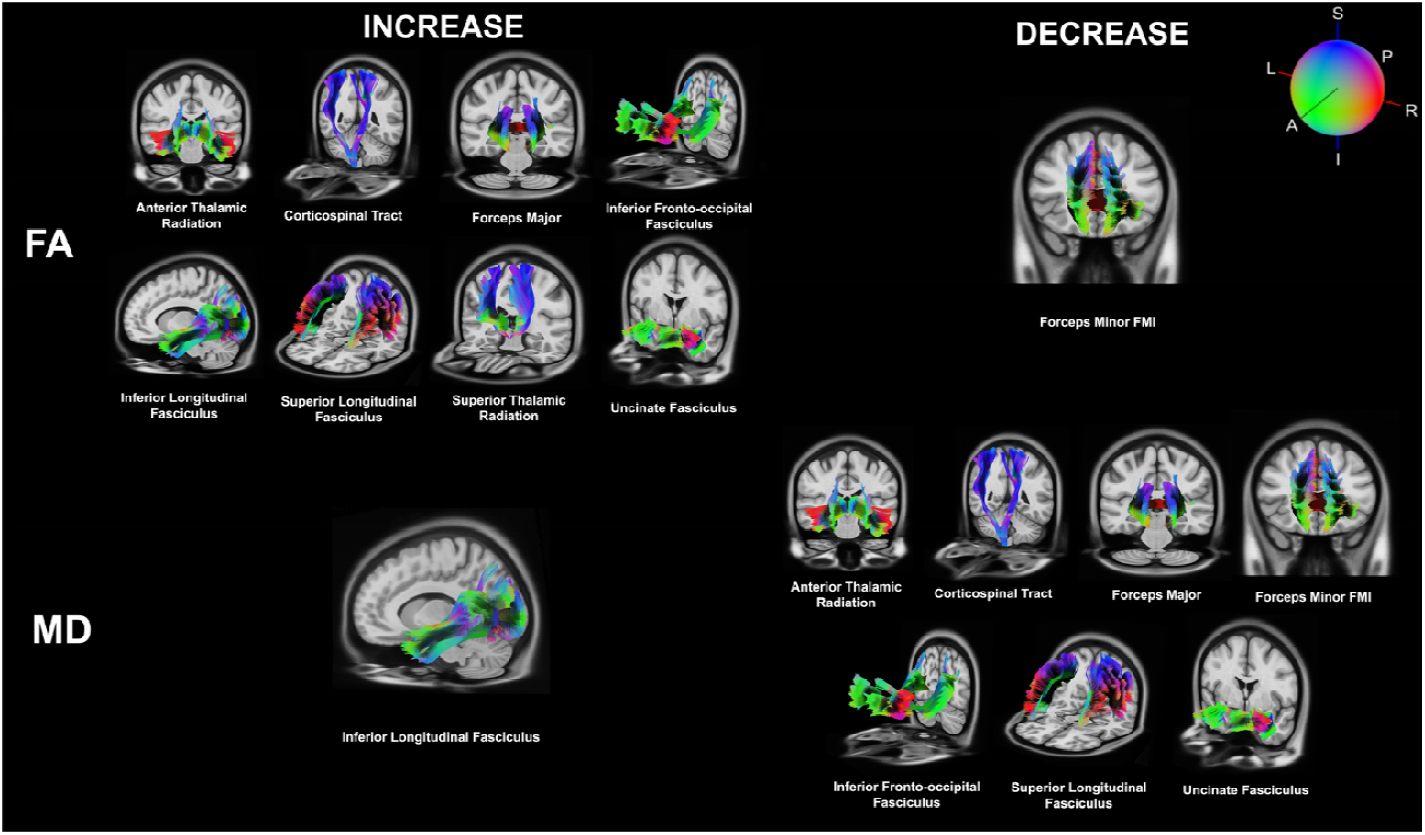
White matter tracts showing significant between-group differences in fractional anisotropy (FA; top) and mean diffusivity (MD; bottom). Tracts with higher FA or MD in the subconcussive exposure group relative to controls are shown on the left, and tracts with lower FA or MD are shown on the right. Streamlines are overlaid on the MNI152 template, and colour indicates principal diffusion direction according to the orientation sphere.

### Statistical Analysis

Data distribution was assessed visually via histograms and Q-Q plots. To examine between-group differences in diffusion metrics at our time points, Welch t-tests were conducted for each WM tract. All statistical analyses was conducted in R (v4.2.2) and Prism (v10). All p-values were adjusted using FDR and statistical significance for all analyses was set at p < 0.05.

Behaviour on the Kinarm BOB task was quantified using a normalized composite measure of overall task performance, the Task Score.^28,29^ The Task Score combines various kinematic metrics such as speed, accuracy, and movement correctness, applying box-cox transformations to yield a normalized value from 0 to 1, where 0 indicates best performance and 1 indicates worst performance.^28,29^ Group differences in the change in Task Score were evaluated using independent t-tests, with Cohen’s d reported as a measure of effect size.

SCAT3 symptom and severity scores were analyzed as change scores (POST-PRE) and compared between groups using Mann-Whitney U tests due to skewing. Glass rank-biserial correlation (r_rb_) was calculated as the effect size for these Mann-Whitney U tests.^30^

## Results

We first wanted to compare if there were any baseline differences at the pre-season timepoint in DTI metrics for each WM tract between our control and subconcussive groups (Figure 1). We found that the subconcussed group had generalized higher FA than our control group in all 14 tracts (all p < 0.001; Figure 2A), except for the forceps minor which had decreased (p < 0.001; Figure 2A). There is also found alongside general lower MD in all tracts (all p < 0.001; Figure 2B) except for left and right interior longitudinal fasciculus which increased (p < 0.001, Figure 2B) and the left and right superior thalamic radiation which had no change (left: p = 0.333, right: 0.915; Figure 2B).

**Figure 2.**
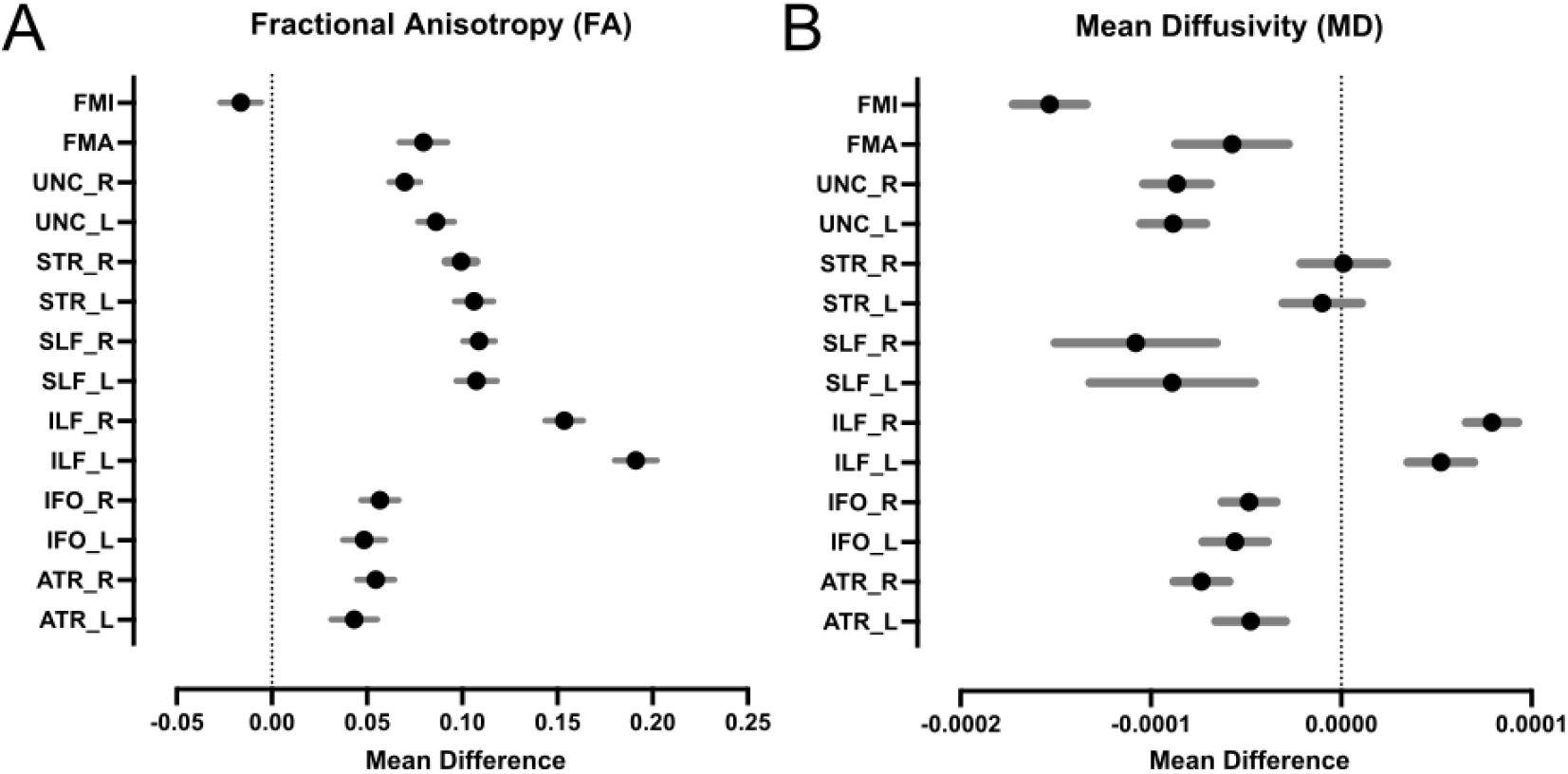
Tract specific diffusion differences at preseason. (A) FA mean differences between contact athletes and non-contact controls across major commissural and association tracts. (B) MD mean differences for the same tracts. Black circles show the mean between group difference for each tract and grey bars show the 95% confidence interval, with the vertical dotted line indicating no difference (mean difference = 0).

Data collected from helmet accelerometers were used to examine the frequency of subconcussive impacts over the football season for each athlete. The 22 athletes enrolled have an average total impacts ranging from 12 to 291 based on position (Table 1). As a sanity check after our separation into two groups based on the median impact frequency, we verified the HE group had received a significantly higher total of impacts than the LE group (p = 0.002, d = 1.02; Table 1). The HE group experienced significantly higher total gForce from these impacts as well (p < 0.001, d = 1.71; Table 1). On average, this was 7.20 impacts per session for the LE group versus 14.62 impacts per session for HE group.

**Table 1.** Total season head-impact exposure by group and playing position. Values are mean ± SD for Total Impacts and Total gForce for high exposure (HE), low exposure (LE), and each position group.

| Variable | HE<br>mean<br>± SD | LE<br>mean<br>± SD | DB<br>(n=6) | WR<br>(n=5) | DL<br>(n=4) | LB<br>(n=3) | OL<br>(n=2) | K<br>(n=1) | RB<br>(n=1) |
| --- | --- | --- | --- | --- | --- | --- | --- | --- | --- |
| Total<br>Impacts | 300<br>± 170 | 154<br>± 94 | 253<br>± 174 | 291<br>± 132 | 281<br>± 133 | 184<br>± 183 | 157<br>± 114 | 12 | 26 |
| Total<br>gForce | 16163<br>± 6486 | 7089<br>± 3799 | 12367<br>± 8510 | 11340<br>± 5113 | 13908<br>± 8637 | 8755<br>± 6377 | 10632<br>± 1576 | 1086 | 20649 |

DB, defensive back; DL, defensive line; K, kicker; LB, linebacker; OL, offensive line; RB, running back; TC, training camp; WR, wide receiver.

Based on the exposure groups described above, we examined bimanual coordination BOB task before and after a single football season. Using the summative task score, we tested whether motor ability on this task changed by the end of the season. Athletes in the LE group showed a modest improvement by the end of the season (−0.33 ± 0.25), whereas those in the HE group became worse over time (0.26 ± 0.33), yielding a between-group difference in change of 0.59 (95% CI 0.28 to 0.89; p < 0.001) and a large effect size (d = 2.07; Figure 3A).

**Figure 3.**
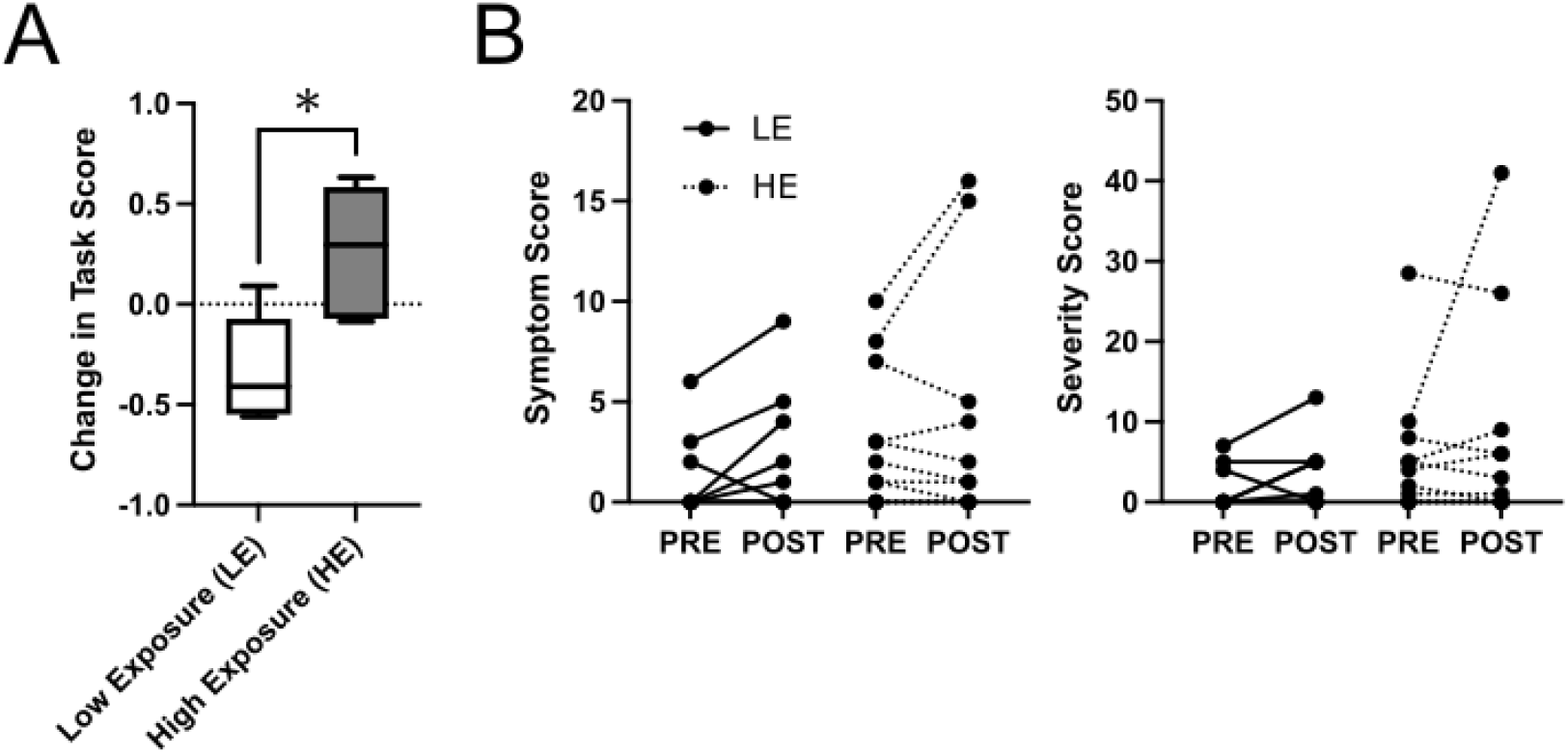
Change in motor task performance and concussion symptoms across the season. (A) Change in Kinarm BOB task score from pre to post season for LE and HE athletes, with asterisk denoting significance. (B) Individual SCAT3 symptom scores (left) and severity scores (right) at PRE and POST season for LE (solid lines) and HE (dashed lines) groups, with lines connecting repeated measures for each athlete.

For the SCAT3, baseline scores were low and comparable between groups, with no significant PRE to POST change within either group (Figure 3B). The change in symptom score did not differ between groups (LE: 1.25 ± 1.91; HE: 0.82 ± 2.93; U = 57, p = 0.29, r_rb_ = 0.30; Figure 3B left). Similarly, the change in severity score did not differ between groups (LE: 1.63 ± 3.42; HE: 2.59 ± 9.62; U = 56, p = 0.33, r_rb_ = 0.27; Figure 3B right). Overall, these findings indicate no group differences in the trajectories of either symptom or severity scores for HE and LE across the season.

## Discussion

RSCIs have increasingly been implicated in subtle yet cumulative alterations in WM microstructure, even in the absence of clinically diagnosed concussion.^2,14^ This study of varsity football players and non-contact controls found that contact athletes began the season with systematically altered white matter microstructure, showing higher FA and predominantly lower MD across most major tracts compared to controls. Over the course of the season, players with higher RSCI exposure, quantified using helmet-mounted accelerometers, developed clear deficits in bimanual coordination on the Kinarm BOB task, whereas LE athletes showed modest improvement. By contrast, SCAT3 symptom and severity scores remained low and did not differ in their trajectories between exposure groups, and longitudinal tract-level DTI changes were minimal. These findings suggest that a history of repetitive subconcussive exposure in contact sport is linked to sustained differences in white matter microstructure and emerging motor control deficits that are not captured by standard symptom-based clinical tools.

### White Matter Microstructural Differences in Contact Sport Athletes

Extensive imaging evidence indicates that athletes exposed to repetitive head impacts show persistent alterations in white matter microstructure. Collision-sport athletes often demonstrate higher FA and lower MD in major white matter tracts compared to non-contact athletes,^31^ although the opposite has also been found.^13^ DTI white matter changes appear even in the absence of any diagnosed concussions, suggesting that years of contact sport participation can induce subtle cumulative changes in brain structure. Consistent with this, Churchill et al. (2017) reported that a greater number of prior sports-related concussions in varsity athletes was associated with increased FA and decreased MD in several major tracts,^32^ a pattern that closely parallels the higher FA and lower MD we observe in athletes with greater subconcussive exposure. Similarly, Kwiatkowski et al. (2024) found decreased MD in the left SLF, IFO, UNC, and ATR, overlapping with the tract-specific MD reductions identified in our cohort.^33^ Longitudinal work further suggests that cumulative head-impact exposure drives progressive DTI change: across both collegiate and high-school cohorts, with the latter exhibiting significantly larger regions of FA increase and MD decrease than controls, indicating that RSCIs induce low-level neurotrauma that accumulates with continued exposure.^34^ Thus confirming that a history of repetitive subconcussive impact is associated with sustained white matter differences, even when no concussions are recorded.

### Subconcussive Impacts and Motor Coordination Deficits

Our results showing declines in bimanual coordination among HE athletes align with an emerging body of work linking repetitive subconcussive hits to subtle motor control impairments. Even a single session of subconcussive head impacts has been shown to measurably alter motor system function. In amateur boxers, sparring without any diagnosed concussions led to increased corticomotor inhibition^35^ and a “dampening” of motor control, evidenced by delayed recruitment of motor units and reduced force output at given activation thresholds.^36^ These findings suggest that repetitive subclinical head trauma can subtly degrade neuromotor integration and coordination. Behaviourally, athletes with greater subconcussive exposure may exhibit slower reaction times, poorer fine motor control, or diminished bimanual task performance, even though such changes might not be obvious without sensitive testing. Although caution is warranted in interpreting these behavioural deficits as task specific, the declining BOB performance in HE athletes, contrasted with improvements in LE athletes, reinforces the idea that accumulating subconcussive blows can impair complex motor tasks. Importantly, such impairments may not be detected by standard clinical symptom scales, underscoring the need for objective motor testing in athletes with higher exposure to head impacts.

### Limitations of Symptom-Based Clinical Assessments

Despite clear group differences in white matter integrity and motor function, we found no corresponding changes in standard clinical concussion evaluations (SCAT3 symptom and severity scores) between HE and LE athletes. Building on Goubran et al. (2023), who found no significant results using the global SCAT score,^37^ we examined SCAT sub-scores to assess symptom and severity changes. Despite white matter alterations, we found no significant differences in sub-scores, reinforcing the concern that SCAT3 may lack the sensitivity to detect subtle, cumulative effects of RSCIs over time. Athletes sustaining repetitive subconcussive hits typically remain asymptomatic longer, so tools relying on self-reported symptoms or brief cognitive exams after exposure register nothing out of the ordinary. The insensitivity of symptom-based assessments to subclinical damage is well-documented; even when neuroimaging or motor testing reveals changes, college athletes frequently show no significant impairments on exams like the SCAT.^31^ Our findings therefore highlight a critical gap in current concussion protocols: conventional sideline or clinical measures may not be sensitive enough to capture effects of cumulative head trauma that falls below the concussion threshold. In this study, players in the HE group objectively declined in a complex motor coordination task, yet their symptom checklists remained unchanged. These results echo the notion that standard concussion assessment tools, while effective for acute injury, lack the sensitivity to detect the subtle neurologic deficits arising from repetitive subconcussive exposure.^31^ More sensitive evaluation methods like advanced imaging, may be required to identify athletes who are experiencing subconcussion-related changes despite feeling fine by subjective report.

## Conclusion

Our findings suggest that a history of repetitive subconcussive impact in contact sport is linked to lasting changes in brain structure and behaviour that routine clinical tools miss. Before the season, varsity football players already showed altered white matter diffusion compared with non-contact controls, and over the season those with higher impact exposure developed clear deficits in bimanual coordination despite negligible symptom scores. These subclinical abnormalities indicate that brief off-season intervals may not be sufficient for full recovery from repeated minor head trauma and may contribute to longer-term risk of cognitive decline and neurodegenerative disease. While the precise timeline and clinical implications remain uncertain, these findings highlight the need for continued efforts to mitigate repetitive head impacts in contact sports. By characterizing how RSCIs influence brain structure and exploring how that structure recovers or degrades over time, researchers and clinicians can better delineate risks, identify thresholds for safe participation, and develop targeted interventions to preserve athlete brain health. The white matter differences and motor deficits highlight the need for proactive monitoring that includes advanced neuroimaging and objective motor assessments, even in athletes who never sustain a diagnosed concussion.

## Data Availability Statement

Data is available upon request.

## References

1. Bailes, J. E., Petraglia, A. L., Omalu, B. I., Nauman, E. & Talavage, T. Role of subconcussion in repetitive mild traumatic brain injury. J. Neurosurg. 119, 1235–1245 (2013).

2. Mainwaring, L., Ferdinand Pennock, K. M., Mylabathula, S. & Alavie, B. Z. Subconcussive head impacts in sport: A systematic review of the evidence. Int. J. Psychophysiol. 132, 39– 54 (2018).

3. Blennow, K., Hardy, J. & Zetterberg, H. The neuropathology and neurobiology of traumatic brain injury. Neuron 76, 886–899 (2012).

4. Broglio, S. P. et al. Head Impacts During High School Football: A Biomechanical Assessment. J. Athl. Train. 44, 342–349 (2009).

5. MacFarlane, M. P. & Glenn, T. C. Neurochemical cascade of concussion. Brain Inj. 29, 139– 153 (2015).

6. Alexander, A. L., Lee, J. E., Lazar, M. & Field, A. S. Diffusion tensor imaging of the brain. Neurother. J. Am. Soc. Exp. Neurother. 4, 316–329 (2007).

7. Tournier, J.-D., Mori, S. & Leemans, A. Diffusion tensor imaging and beyond. Magn. Reson. Med. 65, 1532–1556 (2011).

8. Assaf, Y. & Pasternak, O. Diffusion tensor imaging (DTI)-based white matter mapping in brain research: a review. J. Mol. Neurosci. MN 34, 51–61 (2008).

9. Chun, I. Y. et al. DTI Detection of Longitudinal WM Abnormalities Due to Accumulated Head Impacts. Dev. Neuropsychol. 40, 92–97 (2015).

10. McAllister, T. W. et al. Effect of head impacts on diffusivity measures in a cohort of collegiate contact sport athletes. Neurology 82, 63–69 (2014).

11. Mayer, A. R. et al. A prospective microstructure imaging study in mixed-martial artists using geometric measures and diffusion tensor imaging: methods and findings. Brain Imaging Behav. 11, 698–711 (2017).

12. Caron, B. et al. Advanced mapping of the human white matter microstructure better separates elite sports participation. Preprint at 10.31234/osf.io/dxaqp (2020).

13. Bazarian, J. J. et al. Persistent, Long-term Cerebral White Matter Changes after Sports-Related Repetitive Head Impacts. PLOS ONE 9, e94734 (2014).

14. Bazarian, J. J., Zhu, T., Blyth, B., Borrino, A. & Zhong, J. Subject-specific changes in brain white matter on diffusion tensor imaging after sports-related concussion. Magn. Reson. Imaging 30, 171–180 (2012).

15. Hellewell, S. C., Nguyen, V. P. B., Jayasena, R. N., Welton, T. & Grieve, S. M. Characteristic patterns of white matter tract injury in sport-related concussion: An image based meta-analysis. NeuroImage Clin. 26, 102253 (2020).

16. D’Amario, S. et al. Impact Biomechanics Reveal Positional and Session Type Differences in Canadian Collegiate Football. Eur. J. Sport Sci. 25, e70010 (2025).

17. R Lowrey, C. A Novel Robotic Task for Assessing Impairments in Bimanual Coordination Post-Stroke. Int. J. Phys. Med. Rehabil. s3, (2014).

18. Simmatis, L., Atallah, G., Scott, S. H. & Taylor, S. The feasibility of using robotic technology to quantify sensory, motor, and cognitive impairments associated with ALS. Amyotroph. Lateral Scler. Front. Degener. 20, 43–52 (2019).

19. Montenigro, P. H. et al. Cumulative Head Impact Exposure Predicts Later-Life Depression, Apathy, Executive Dysfunction, and Cognitive Impairment in Former High School and College Football Players. J. Neurotrauma 34, 328–340 (2017).

20. Kelley, A. M., Bernhardt, K., Hass, N. & Rooks, T. Detecting functional deficits following sub-concussive head impacts: the relationship between head impact kinematics and visual-vestibular balance performance. Brain Inj. 35, 812–820 (2021).

21. Haran, F. J., Handy, J. D., Servatius, R. J., Rhea, C. K. & Tsao, J. W. Acute neurocognitive deficits in active duty service members following subconcussive blast exposure. Appl. Neuropsychol. Adult 28, 297–309 (2021).

22. Guskiewicz, K. M. et al. Evidence-based approach to revising the SCAT2: introducing the SCAT3. https://doi.org/10.1136/bjsports-2013-092225 (2013) xdoi:10.1136/bjsports-2013-092225.

23. Slobounov, S. M. et al. The effect of repetitive subconcussive collisions on brain integrity in collegiate football players over a single football season: A multi-modal neuroimaging study. NeuroImage Clin. 14, 708–718 (2017).

24. Champagne, A. A., Coverdale, N. S., Germuska, M., Bhogal, A. A. & Cook, D. J. Changes in volumetric and metabolic parameters relate to differences in exposure to sub-concussive head impacts. J. Cereb. Blood Flow Metab. Off. J. Int. Soc. Cereb. Blood Flow Metab. 40, 1453–1467 (2020).

25. Jenkinson, M., Beckmann, C. F., Behrens, T. E. J., Woolrich, M. W. & Smith, S. M. FSL. NeuroImage 62, 782–790 (2012).

26. Behrens, T. E. J., Berg, H. J., Jbabdi, S., Rushworth, M. F. S. & Woolrich, M. W. Probabilistic diffusion tractography with multiple fibre orientations: What can we gain? NeuroImage 34, 144–155 (2007).

27. Warrington, S. et al. XTRACT - Standardised protocols for automated tractography in the human and macaque brain. NeuroImage 217, 116923 (2020).

28. Simmatis, L. E. R., Early, S., Moore, K. D., Appaqaq, S. & Scott, S. H. Statistical measures of motor, sensory and cognitive performance across repeated robot-based testing. J. NeuroEngineering Rehabil. 17, 86 (2020).

29. Scott, S. H., Lowrey, C. R., Brown, I. E. & Dukelow, S. P. Assessment of Neurological Impairment and Recovery Using Statistical Models of Neurologically Healthy Behavior. Neurorehabil. Neural Repair 37, 394–408 (2023).

30. Tomczak, M. & Tomczak, E. The need to report effect size estimates revisited. An overview of some recommended measures of effect size. 1, (2014).

31. Churchill, N. W., Hutchison, M. G., Di Battista, A. P., Graham, S. J. & Schweizer, T. A. Structural, Functional, and Metabolic Brain Markers Differentiate Collision versus Contact and Non-Contact Athletes. Front. Neurol. 8, (2017).

32. Churchill, N. et al. Brain Structure and Function Associated with a History of Sport Concussion: A Multi-Modal Magnetic Resonance Imaging Study. J. Neurotrauma 34, 765– 771 (2017).

33. Kwiatkowski, A. et al. Uncovering the hidden effects of repetitive subconcussive head impact exposure: A mega-analytic approach characterizing seasonal brain microstructural changes in contact and collision sports athletes. Hum. Brain Mapp. 45, e26811 (2024).

34. Jang, I. et al. Every hit matters: White matter diffusivity changes in high school football athletes are correlated with repetitive head acceleration event exposure. NeuroImage Clin. 24, 101930 (2019).

35. Di Virgilio, T. G. et al. Evidence for Acute Electrophysiological and Cognitive Changes Following Routine Soccer Heading. EBioMedicine 13, 66–71 (2016).

36. Di Virgilio, T. G., Ietswaart, M., Wilson, L., Donaldson, D. I. & Hunter, A. M. Understanding the Consequences of Repetitive Subconcussive Head Impacts in Sport: Brain Changes and Dampened Motor Control Are Seen After Boxing Practice. Front. Hum. Neurosci. 13, (2019).

37. Goubran, M. et al. Microstructural Alterations in Tract Development in College Football and Volleyball Players: A Longitudinal Diffusion MRI Study. Neurology 101, (2023).

